# A dual-color miniature endoscope for calcium imaging in behaving mice

**DOI:** 10.64898/2026.02.19.706914

**Authors:** Jinyong Zhang, Feiyang Hong, Jiwon Kim, Konstantin Bakhurin, Namsoo Kim, Henry H. Yin

## Abstract

Calcium imaging with miniature endoscopes has become an essential tool in neuroscience, but conventional miniscopes typically record signals from only a single calcium indicator. Here, we present a dual-color miniature endoscope (miniscope) that enables simultaneous calcium imaging from two neuronal populations using spectrally distinct genetically encoded indicators. In freely moving mice, we used this system to record activity from striatal neurons of the direct (dSPN) and indirect (iSPN) pathways. We showed that dSPNs were activated earlier than iSPNs during contraversive movements, with dSPNs preferentially active during acceleration and iSPNs during deceleration. During ipsiversive turns, however, this temporal relationship was reversed. These findings indicate that dSPNs and iSPNs are not concurrently active, but instead exhibit complementary, direction-dependent dynamics that govern movement velocity. Our dual-color miniscope provides a compact, cost-effective platform for simultaneous two-population imaging, offering new opportunities to dissect coordinated activity across neural circuits in freely behaving animals.

## Introduction

Calcium imaging has become an essential tool in neuroscience research. The development of microscopes for imaging fluorescent signals have enabled researchers to visualize neural signals in awake, behaving animals^1–5^. The growing availability of genetically encoded calcium indicators (GECIs) enables simultaneous imaging of multiple neural populations. However, current imaging approaches impose substantial limitations on dual-color calcium imaging in freely moving animals.

Two-photon (2P) microscopes offer excellent spatial resolution and availability of dual-color imaging^6^, but they have traditionally been limited to head-fixed preparations^7^. Only recently have miniature two-photon microscopes been developed for use in freely moving animals, but they remain costly due to their demanding light source requirements^1,8,9^. In contrast, fiber photometry readily enables dual-color recording through distinct calcium indicators but suffers from limited spatial resolution, which makes it impossible to record activity from individual neurons^10,11^.

One-photon miniature fluorescence microscopes offer a valuable alternative, providing cellular-level resolution while being compact enough for unrestricted movement in freely moving animals^2,4,12^. However, traditional setups typically allow imaging from only one channel at a time. Here, we introduce a dual-color miniscope designed for calcium imaging in behaving mice. This miniscope uses a dichroic mirror in the emission path, splitting fluorescence signals onto two independent image sensors. Two light-emitting diodes (LEDs) with different wavelengths (blue, 450–490 nm; and lime, 547–572 nm) and two Complementary Metal-Oxide-Semiconductor (CMOS) sensors are used to record two channels simultaneously with minimal crosstalk.

Using this dual-color miniscope, we recorded activity from two distinct populations of striatal projection neurons (SPNs). The striatum is the main input nucleus of the basal ganglia^13–15^. Most neurons in the striatum are SPNs, comprising two distinct populations that are spatially intermixed^16^: striatonigral neurons (direct pathway SPNs or dSPNs), expressing D1-like dopamine receptors, and striatopallidal (indirect pathway SPNs or iSPNs) expressing D2 receptors and A2A adenosine receptors. These two neural populations are believed to play complementary roles in behavior ^13,17–20^. Previous work showed that both dSPNs and iSPNs are activated during volitional movements^21–23^, although their temporal and functional relationships are not fully understood.

Using two distinct calcium indicators (Cre-on jRCaMP1b and Cre-off GCaMP6s), we recorded activity from both dSPNs and iSPNs simultaneously using our dual-color miniscope. Our results demonstrate that dSPNs and iSPNs are not concurrently activated, contrary to previous findings^21^. Instead, dSPN activity typically precedes iSPN activity in contraversive movement (towards the side contralateral to the recording side), with dSPNs preferentially activated during acceleration and iSPNs activated during deceleration. Our dual-color miniscope is therefore capable of revealing subtle temporal differences in the timing of activity from defined neuronal populations, thereby improving our understanding of basal ganglia function.

## Results

### System design

A dual-color miniscope was developed based on the UCLA V3 Miniscope platform (including the CMOS sensor, DAQ, and software). The housing was redesigned to enable simultaneous imaging of GCaMP and RCaMP, though the system can be readily adapted to visualize any two spectrally distinct fluorescent indicators by selecting the appropriate LEDs and optical filters. Two LEDs are positioned at distinct locations for fluorescence excitation: a blue LED (450–490 nm filter) for GCaMP calcium indicator excitation and a lime-green LED (547–572 nm filter) for jRCaMP excitation. The blue LED is driven by a constant-current power source delivering up to 30 mA in 1 mA increments, while the lime-green LED is controlled by a separate power source providing up to 200 mA in 4 mA increments. Both LEDs achieve a maximum intensity of 2 mW/mm², measured beneath a gradient-index (GRIN) lens. The two light wavelengths are combined in the excitation path using an excitation dichroic mirror (T510lpxr).

Fluorescence emitted by the two indicators is separated by an emission dichroic mirror, directing light to two distinct CMOS sensors. An achromatic lens in the main emission path ensures proper focus onto the CMOS sensors (Figure 1). To mitigate chromatic aberration from the GRIN lens, each CMOS sensor is mounted on a sliding sleeve, allowing focal plane adjustments (Supplementary Fig. 2B) ^3^. All the materials used are listed in **Table 1**.

**Figure 1.**
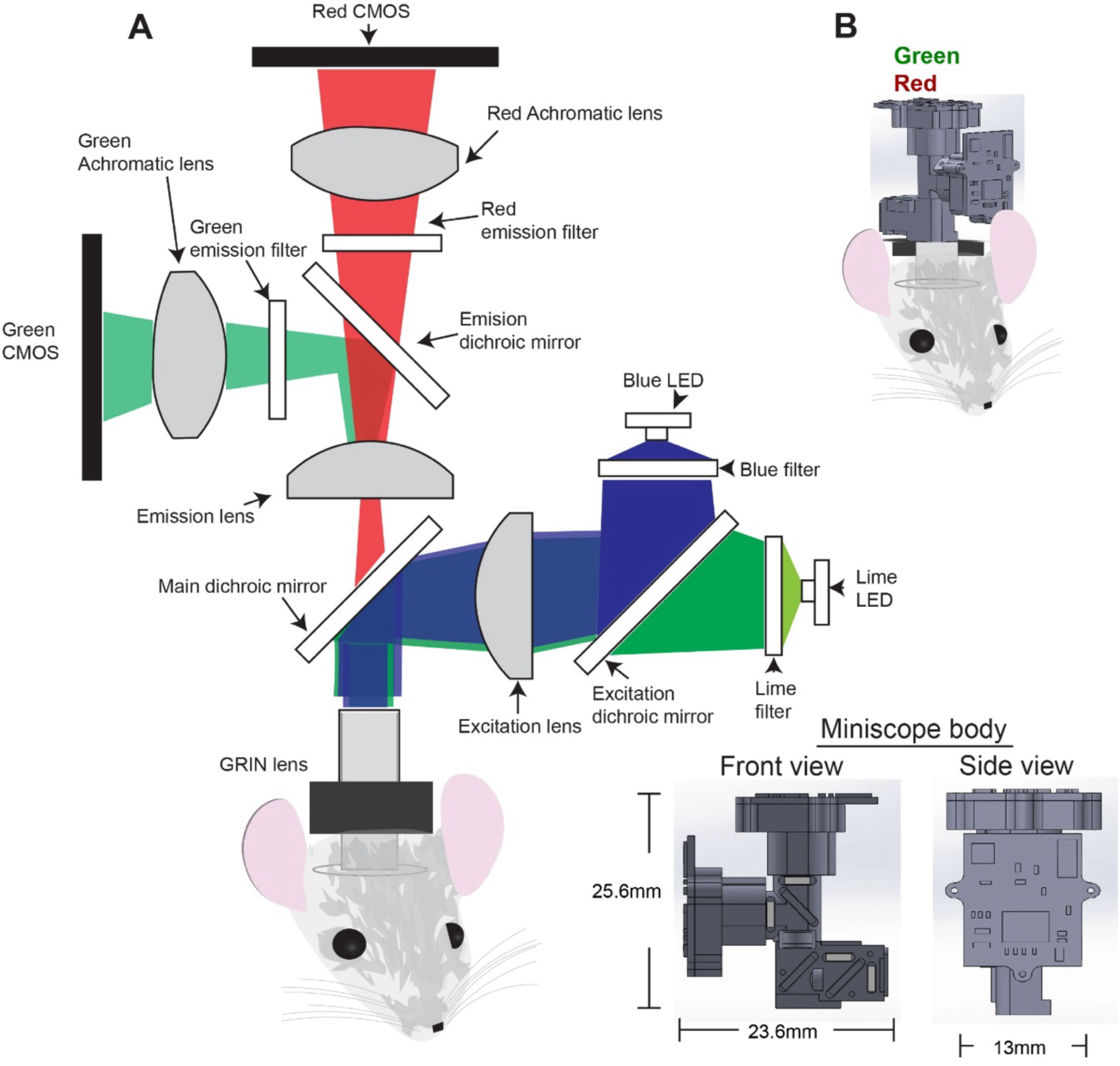
Schematic of Dual-color Miniscope. A) Schematic of the dual-color miniscope components. Two LED light sources are used in the excitation path: a blue LED for GCaMP6s excitation and a lime LED for jRCaMP1b. The emission path is split into two channels, each detected by an independent CMOS sensor—one dedicated to green light (GCaMP6s) and the other to red light (jRCaMP1b). The CMOS sensors are positioned at focal planes matched by achromatic lenses. B) Illustration of the dual-color miniscope mounted on a mouse head.

**Table 1.** Parts for dual-color miniscope.

| Item | Manufacturer | Manufacturer P/N | Description | Quantity |
| --- | --- | --- | --- | --- |
| Lime LED | LUXEON | LXML-PX02-0000 | 566-569nm LED | 1 |
| Blue LED | LUXEON | LXML-PB01-0040 | 450-490nm LED | 1 |
| Lime filter | Chroma | ET560/25x | Lime excitation filter 547.5nm-572.5nm, 4×4×1mm | 1 |
| Blue filter | Chroma | ET470/40x | Blue excitation filter 450nm-490nm, 4×4×1mm | 1 |
| Excitation dichroic mirror | Chroma | T510lpxr | Excitation dichroic mirror 4×6×1mm | 1 |
| Excitation lens | Edmund optics | 45141 | 3.0mm Dia. x 4.5mm FL, Plane-Convex lens | 1 |
| Emission lens | Edmund optics | 45408 | 5.0mm Dia. x 20mm FL, Achromatic lens | 1 |
| GRIN lens | Edmund optics | 64519 | 1.8mm Dia, 670nm DWL, 0.0mm WD | 1 |
| Main dichroic mirror | Chroma | 69013bs | Main dichroic mirror 4×6×1mm | 1 |
| Emission dichroic mirror | Chroma | T550lpxrt | Emission dichroic mirror 4×6×1mm | 1 |
| Green emission filter | Chroma | ET525/36m | Green emission filter 4×4×1mm | 1 |
| Green Achromatic lens | Edmund optics | 45206 | 5.0mm Dia. x 10mm FL, Achromatic lens | 1 |
| Red emission filter | Chroma | ET630/75m | Red emission filter 4×4×1mm | 1 |
| Red Achromatic lens | Edmund optics | 45206 | 5.0mm Dia. x 10mm FL, Achromatic lens | 1 |
| Green CMOS/ Red CMOS | UCLA miniscope | UCLA miniscope V3 | CMOS, DAQ and software | 2 |

The dual-color miniscope weighs 6.6 g. It provides a resolution of 608 × 608 pixels, a field of view of 500 × 500 µm, and an overall magnification of 20x (**Table 2**). The housing is fabricated using Multi-Jet Fusion 3D printing with Nylon PA12, ensuring high durability and low weight. Because the two LEDs, filters, and CMOS sensors are independently positioned, the system is highly modular and adaptable for imaging various combinations of indicators. By replacing the LEDs, excitation and emission filters, and dichroic mirrors, the optical configuration can be readily adjusted to match the specific spectral properties of the chosen indicators. The system is also cost-effective (parts listed in Table 1).

**Table 2.** Comparisons of different one-photon Dual-color miniscopes currently available.

|  | Weight(g) | Size (L by W by H, mm) | Number of CMOS sensors | Number of LEDs | Number of channels | Maximum frame rate(fps) | Source |
| --- | --- | --- | --- | --- | --- | --- | --- |
| Dual-color miniscope | 6.6 | 23.6 by 13 by 25.6 | 2 | 2 | 2 | 60 (each channel) | This paper |
| Dual-channel miniscope | 4.8 | 13.96 by 11.32 by 31.55 | 2 | 1 | 2 | 30(each channel) | Open source |
| Doric lenses 2-color | 3.7 | 17 by 9.5 by 18 | 1 | 1 | 2 | 45 | Commercial |
| Inscopix NVue | 2.4 | 14.8 by 8.5 by 21.5 | 1 | 1 | 2 | 30 | Commercial |

### Crosstalk testing

To evaluate potential crosstalk between the excitation LEDs and fluorescence detection channels, the miniscope was placed inside a light-tight black box, and images were acquired while each LED was activated separately at maximum power. No change in brightness was observed when the blue LED was turned on. To assess crosstalk between the green (GCaMP6s) and red (jRCaMP1b) channels, we performed imaging experiments with a single calcium indicator^24^. Using a Cre-dependent GCaMP6s virus injected into D1-Cre mice (Figure 2B), we observed strong fluorescence signals from the blue channel (Figure 2C). On the other hand, even at the maximum blue LED power (30 mA), fluorescence in the red channel was minimal. In addition, we injected a Cre-dependent jRCaMP1b virus into D1-Cre mice, and strong fluorescence was observed in the red channel at maximum lime LED power, with low signal in the blue channel (Figure 2B, C). When RCaMP-expressing mice were exposed to blue LED light and GCaMP-expressing mice to lime LED light, fluorescence in both channels remained minimal (Figure 2C).

**Figure 2.**
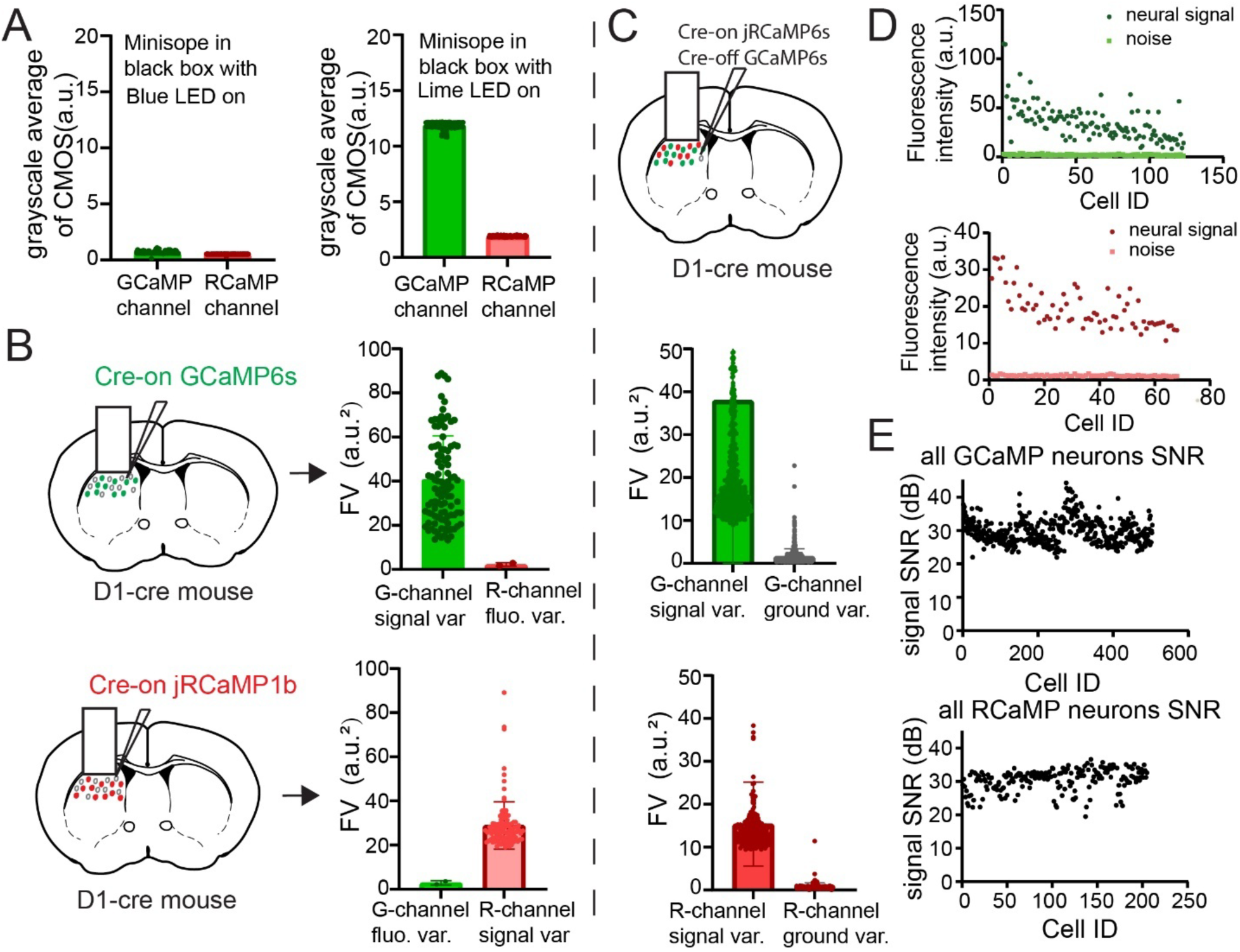
Crosstalk between GCaMP6s and jRCaMP1b channels. A) Test of two types of LED light impact on CMOS. Miniscope was placed in a light-tight box with the blue LED operating at maximum power. The average grayscale value was calculated for each frame across 1,000 frames (left). The average grayscale value was computed for each of the 1,000 frames while the lime LED operated at maximum power (right). B) D1-Cre mice expressing exclusively Cre-dependent GCaMP6s(n=2) or jRCaMP1b(n=2) (left). The signal/fluorescence variance of two channels with both LED functioning properly (right). C) Top, D1-Cre mice expressing both GCaMP6s and jRCaMP1b (n=5). Fluorescence variance of neuronal fluorescence compared with background in GCaMP6s channel (n=5, middle) and jRCaMP1b channel (n=5, bottom). D) Fluorescence intensity compared with background from a representative mouse in GCaMP6s channel(top) and jRCaMP1b(bottom). E) Signal-to-noise (SNR) in co-expressed GCaMP6s and jRCaMP1b mice (n=5), GCaMP channel (top, 504 cells) and jRCaMP1b channel (bottom, 205 cells).

The LUXEON Rebel lime LED (nominal peak wavelength 566–569 nm) exhibits a broad emission spectrum, and despite the presence of an excitation filter (ET560/25x), a small fraction of its light overlaps with the GCaMP emission band. This consistent, wavelength-dependent overlap produced minor crosstalk. To correct for LED crosstalk, we implemented pixel-by-pixel flat-field correction. We acquired full-field crosstalk images by placing the miniscope in a light-tight box and imaging with each LED at maximum power separately (without any fluorescent sample). These images capture the spatial pattern of LED leakage across the entire field of view, including effects such as vignetting or uneven illumination distribution. This approach accounts for spatial non-uniformities and provides more rigorous and spatially accurate crosstalk correction.

### Fluorescence crosstalk in exclusively expressed mice

To evaluate fluorescence-based crosstalk between the green (GCaMP6s) and red (jRCaMP1b) channels, we conducted imaging experiments in mice expressing only one calcium indicator at a time. For GCaMP6s imaging, a Cre-dependent GCaMP6s virus was injected into D1-Cre mice (n = 2; Figure 2B). Under both blue and lime LED illumination, the GCaMP channel exhibited strong neuronal fluorescence, with 92% of neuronal fluorescence variance exceeding 20, indicating high signal discriminability. In contrast, no detectable fluorescence was observed in the red channel, and the variance of frame-averaged brightness remained below 3. For jRCaMP1b imaging, a Cre-dependent jRCaMP1b virus was injected into D1-Cre mice (n = 2; Figure 2B). The jRCaMP1b channel displayed robust fluorescence, with all neuronal signal variances exceeding 20, while the GCaMP channel showed minimal crosstalk (frame-averaged brightness variance < 3).

We then used the dual-color miniscope to record SPNs co-expressing GCaMP6s and jRCaMP1b (Figures 3A and 4A). Fluorescence signals were simultaneously captured from the green (450–490 nm, GCaMP6s) and red (547–572 nm, jRCaMP1b) channels under concurrent blue and lime LED excitation (Figure 1). The miniscope was mounted on a chronically implanted GRIN lens (1.8 mm diameter, 4.3 mm length) in freely moving mice.

**Figure 3.**
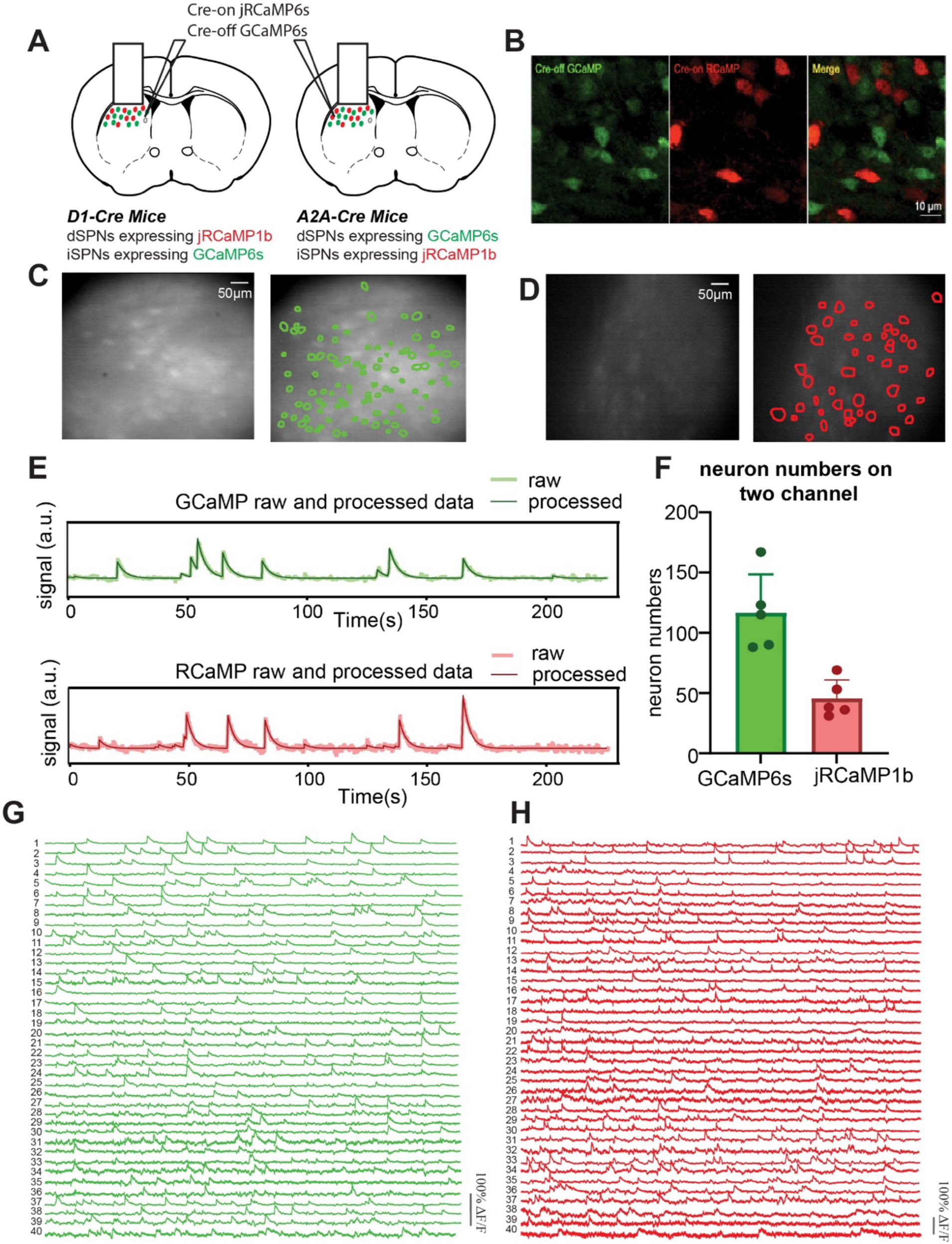
Dual-color calcium imaging from the dorsal striatum in freely moving mice. A) Co-expression of GCaMP6s and jRCaMP1b in striatal SPNs. Both D1-Cre and A2A-Cre mouse lines were used in the experiments. B) Labeling of dSPNs and iSPNs. C) Temporal averaged image of the GCaMP6s channel from one mouse (Left), and ROI contours extracted using the CNMF algorithm overlaid on the temporal average image (Right). D) jRCaMP1b channel (Left) and corresponding ROI contours from the same mouse as in panel C (Right). E) Example traces from both channels, showing raw (light color) and deconvolved (dark color) signals. F) Number of neurons extracted from each channel using CNMF. G) Example raw traces from the GCaMP6s channel. H) Example raw traces from the jRCaMP1b channel.

**Figure 4.**
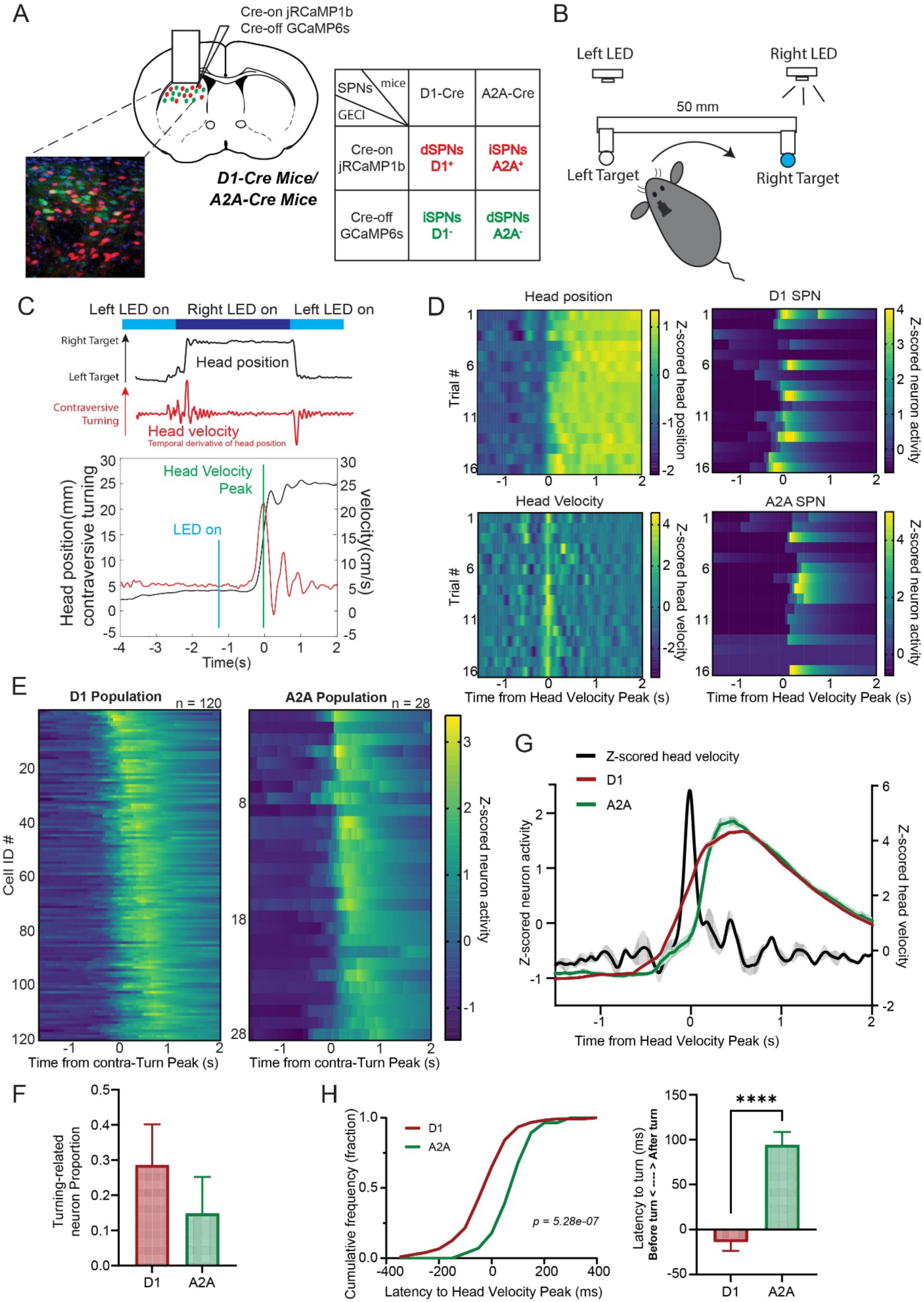
Coordination of dSPNs and iSPNs during contraversive turning in freely moving mice. A) Dual-color miniscope calcium imaging of dSPNs and iSPNs in the dorsal striatum. B) Schematic of the switching task. Mice moved between two water spouts positioned 50 mm apart on a behavioral platform. An LED positioned above each spout served as a visual cue indicating the currently active water source. When the LED is turned on, it indicates that the spout next to the LED has water. Thus the mouse has to switch between different target spouts to obtain water. Kinematics was recorded using a 3D motion capture system. C) Representative event from a session in which neural activity was recorded in the left hemisphere. Head position along the left–right water spout axis was continuously tracked as the reward and cue LED switched to the contralateral side. Head velocity was calculated as the temporal derivative of head position. D) Behavioral description and calcium signals from a representative session. One direct pathway SPN and one indirect pathway SPN are shown. E) Average calcium signals of selected contraversive turning-related dSPNs (Left, n=2, cells=120) and iSPNs (Right, n=2, cells=28), aligned to the head velocity peak and sorted by peak activity time. We recorded a total of 262 dSPNs and 201 iSPNs from 2 D1-Cre and 2 A2A-Cre mice. A2A-Cre mice have 230 dSPNs and 52 iSPNs, whereas D1-Cre mice have 33 dSPNs and 149 iSPNs. F) Proportion of contraversive turning-related neurons. G) Z-score of dSPNs activity (red, left y-axis), iSPNs activity (green, left y-axis) and velocity (black, right y-axis) aligned to the head velocity peak. The above SPNs are the same populations as shown in E). H) Time to head velocity peak for dSPNs (red) and iSPNs (green) during contraversive turning. Left, cumulative distribution. A two-sample Kolmogorov–Smirnov test revealed a significant difference between the two distributions (p = 5.28 × 10⁻⁷). Right, average time to peak velocity. **** p < 0.0001, two-tailed unpaired t-test.

Under both excitation conditions, both channels exhibited robust neuronal fluorescence signals. For GCaMP imaging, over 99% of neuronal fluorescence variance exceeded 10, compared to an average background variance of 1.58. In the jRCaMP1b channel, all neuronal fluorescence variances exceeded 10, with an average background variance of 0.26 (n = 5; Figure 2C). In a representative mouse (Figure 2D), the average fluorescence intensity of neuronal signals was 35.23 for the GCaMP6s channel (123 neurons) and 19.29 for the jRCaMP1b channel (68 neurons). Across five animals, all GCaMP6s-labeled neurons (n = 608) and 99% of jRCaMP1b-labeled neurons (n = 205) exhibited >20 SNR (Figure 2E).

Both GCaMP and RCaMP viruses were clearly expressed in the same region (Figure 3B). However, due to competition between the two viral constructs, the jRCaMP1b signal appeared dimmer in co-expression experiments (average fluorescence variance = 15.36) compared with single-expression experiments (average variance = 28.89). We applied a post-processing enhancement step to the RCaMP channel prior to neuron trace extraction (Supplementary Video 2). Example traces from the same mouse showed clearly independent activity patterns in both channels (Figure 3E). Across five mice, the GCaMP6s channel yielded an average of 106 neuron traces, while the jRCaMP1b channel yielded 48 traces (Figure 3F).

### Dual-Color imaging

We used dual-color calcium imaging in striatal SPNs co-expressing GCaMP6s and jRCaMP1b (Figures 3A, 4A). During simultaneous blue (450–490 nm) and lime (547–572 nm) illumination, neuronal activity was recorded concurrently in the green (GCaMP6s) and red (jRCaMP1b) channels (Figure 1).

A chronically implanted GRIN lens (1.8 mm diameter, 4.3 mm length) was used. Both GCaMP6s and jRCaMP1b were co-expressed within the same striatal region (Figure 3B). However, during dynamic calcium imaging, the jRCaMP1b signal appeared dimmer due to competitive expression with GCaMP6s, yielding an average brightness of ∼40 gray levels (8-bit scale). To improve visualization, we applied a uniform enhancement step to the jRCaMP1b videos prior to single-neuron analysis (Supplementary Video 2). In a representative mouse, we identified 150 neurons in the GCaMP6s channel and 68 neurons in the jRCaMP1b channel (Figure 3G–H, Supplementary Video 1). After applying a fixed baseline correction for LED-induced crosstalk (see Methods), the signal-to-noise ratio (SNR) was computed for each neuron. Across four animals, 95% of GCaMP6s-labeled neurons (n = 500) and 99% of jRCaMP1b-labeled neurons (n = 180) exhibited SNR > 20 (Figure 2D–E).

### Activity of dSPNs and iSPNs in relation to behavior

To simultaneously record neural activity in striatal dSPNs and iSPNs, we co-expressed Cre-dependent RCaMP1b (Flex-jRCaMP1b) and Cre-independent GCaMP6s (DO(FAS)-GCaMP6s) in the striatum of D1-Cre and A2A-Cre mice (Figure 3A). In this preparation, dSPNs in D1-Cre mice and iSPNs in A2A-Cre mice expressed RCaMP1b, whereas putative dSPNs in A2A-Cre mice and putative iSPNs in D1-Cre mice expressed GCaMP6s (Figure 4A). Immunostaining confirmed distinct expression of the two sensors in separate neuronal populations (Figure 3B).

Neural activity was recorded using a chronically implanted gradient-index (GRIN) lens (1.8 mm diameter, 4.3 mm length). Both blue and lime LEDs were illuminated to enable simultaneous calcium imaging of both GCaMP and RCaMP channels. Each of the two CMOS sensors was connected to a dedicated UCLA DAQ system, so that the imaging data was recorded along with the frame timestamps (Supplementary Figure 1).

After recording, frame signals were synchronized to align data from both channels. If any frame loss was detected (which was exceedingly rare), the missing frame was replaced by duplicating the adjacent frame, thereby preserving temporal alignment across both imaging channels and with other experimental events (Supplementary Figure 2).

Neural activity was extracted from the fluorescence traces of each unit and deconvolved using a modified constrained nonnegative matrix factorization (CNMF) algorithm (Figure 3C, D)⁴². Striatal dSPNs and iSPNs were simultaneously imaged over a 3-minute period at 20 frames per second (Figure 3). In our best mouse, we recorded 180 neurons from the GCaMP6s channel and 68 neurons from the RCaMP1b channel within a 600 × 600 µm field of view (FOV). Fluorescence traces from the two channels did not overlap, suggesting minimal or no crosstalk (Figure 3F).

### Direct and Indirect pathway activity during behavior

Water-restrained mice were placed on a platform with two water spouts positioned 50 mm apart, each paired with an LED indicator. During the task, 12 uL 20% sucrose droplets were delivered every 600 ms from one of the two spouts. The spouts alternated in delivering water rewards, with the active spout indicated by a corresponding LED (Figure 4B). Mice made contraversive and ipsiversive turns to switch between the two spouts throughout the session. The relative position of the mouse head to the two waterspouts was continuously tracked. Using the axis defined by the line connecting the two spouts, head movements were quantified as changes in position along this axis, and head velocity was subsequently calculated based on head position data (Fig. 4C).

Dual-color calcium imaging observed a number of both direct and indirect pathway SPNs whose activation is associated with contraversive turning (Fig. 4D). Turning-related SPNs were defined as those activated during more than half of the turning events within a one-second window around the head velocity peak (Supplementary Figure 3). Following this criterion, 120 D1^+^ SPNs and 28 A2A^+^ SPNs were identified (Fig. 4E). The activity of dSPNs preceded that of iSPNs by 110 ±22 ms (Figure 4H). Many dSPNs are activated during acceleration (just before the velocity peak) and iSPNs were more commonly activated during deceleration (just after the velocity peak) during contraversive turning (Figure 4G).

### Load cell experiments

To further confirm the timing of dSPN and iSPN activity, we used a head-fixation setup to quantify direction-specific movements using force sensors^25,26^. This multiple-load-cell head-fixed device enabled measurement of force along orthogonal axes of movement (up/down, left/right, forward/backward) (Figure 5A). Force measured over time is the rough equivalent of velocity, but load cells allow monitoring of the movement with extremely high sensitivity and temporal precision^25–28^.

**Figure 5.**
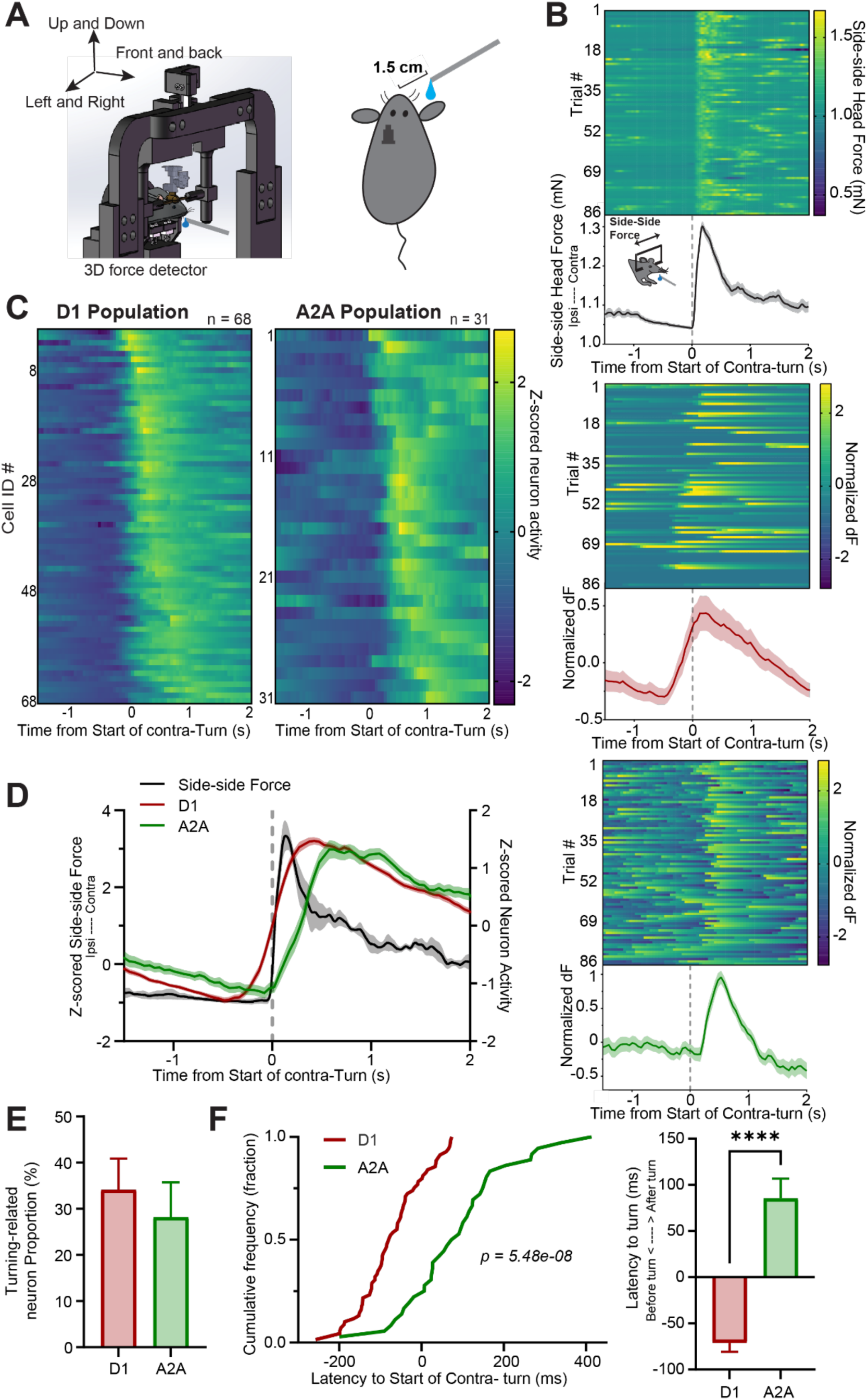
Coordination of dSPNs and iSPNs during contraversive force exertion in head-fixed mice. A) Head-fixation platform with 3D load cell force recording (1,000 Hz) and simultaneous dual-color miniscope imaging (20 fps). The water spout was positioned contralateral to the recorded hemisphere. B) Representative session showing contraversive force exertion (Top), with corresponding activity of example D1 SPNs (Middle) and A2A SPNs (Bottom). C) Average calcium signals of all contraversive pushing-related dSPNs (Left, n=2, cells=68) and iSPNs (Right, n=2, cells=31), aligned to the start of force exertion and sorted by peak activity time (We recorded a total of 229 dSPNs and 128 iSPNs from 2 D1-Cre and 2 A2A-Cre mice. Specifically, A2A-Cre mice contributed 191 dSPNs and 32 iSPNs, whereas D1-Cre mice contributed 38 dSPNs and 96 iSPNs). D) Z-score of dSPNs activity (red, right y-axis),iSPNs activity (green, right y-axis) and side-side force (black, left y-axis) aligned to the start of force exertion. The above SPNs are the same populations as shown in C). E) Proportion of contraversive pushing-related neurons. F) Time to the start of force exertion for dSPNs (red) and iSPNs (green). (Left) Cumulative distribution plots. A two-sample Kolmogorov–Smirnov test revealed a significant difference between the two distributions (p = 5.48 × 10⁻^8^). Right, average time to push. **** p < 0.0001, two-tailed unpaired t-test.

The head of the mouse was fixed using a metallic head bar, and turning-like behavior in three dimensions could be reliably detected by monitoring changes in load cell voltage. A water spout was placed 1.5 cm from the mouse’s mouth, allowing the mouse to obtain 10% sucrose by making a large lateral push similar to a turning movement (Figure 5A). After behavioral training, the activity of striatal dSPNs and iSPNs was simultaneously recorded. When the waterspout was positioned contralateral to the recorded hemisphere, contraversive movement-related neurons were identified using the same criteria as in the freely moving turning task (Supplementary Figure 4). Neurons that were activated during more than half of the lateral movement trials were classified as turning-related (Figure 5B). Using this criterion, 68 D1+ SPNs and 31 A2A+ SPNs were identified (Figure 5C, 5E). Consistent with earlier findings (Figure 4), dSPN activity preceded iSPN activity by ∼160 ms (Figure 5D), with dSPNs leading the onset of force exertion by 71 ms and iSPNs lagging by 86 ms on average (Figure 5F).

Notably, when the waterspout was placed ipsilateral to the recorded hemisphere, ipsiversive turning-related SPNs were identified in the same group of mice (Figure 6A), though in smaller numbers: 22 D1+ SPNs and 12 A2A+ SPNs (Figure 6B, C). In contrast to contraversive movements, iSPNs were activated prior to dSPNs during ipsiversive movements (Figure 6D), with iSPNs leading the onset of force exertion by 61 ms and dSPNs lagging by 58 ms (Figure 6E).

**Figure 6.**
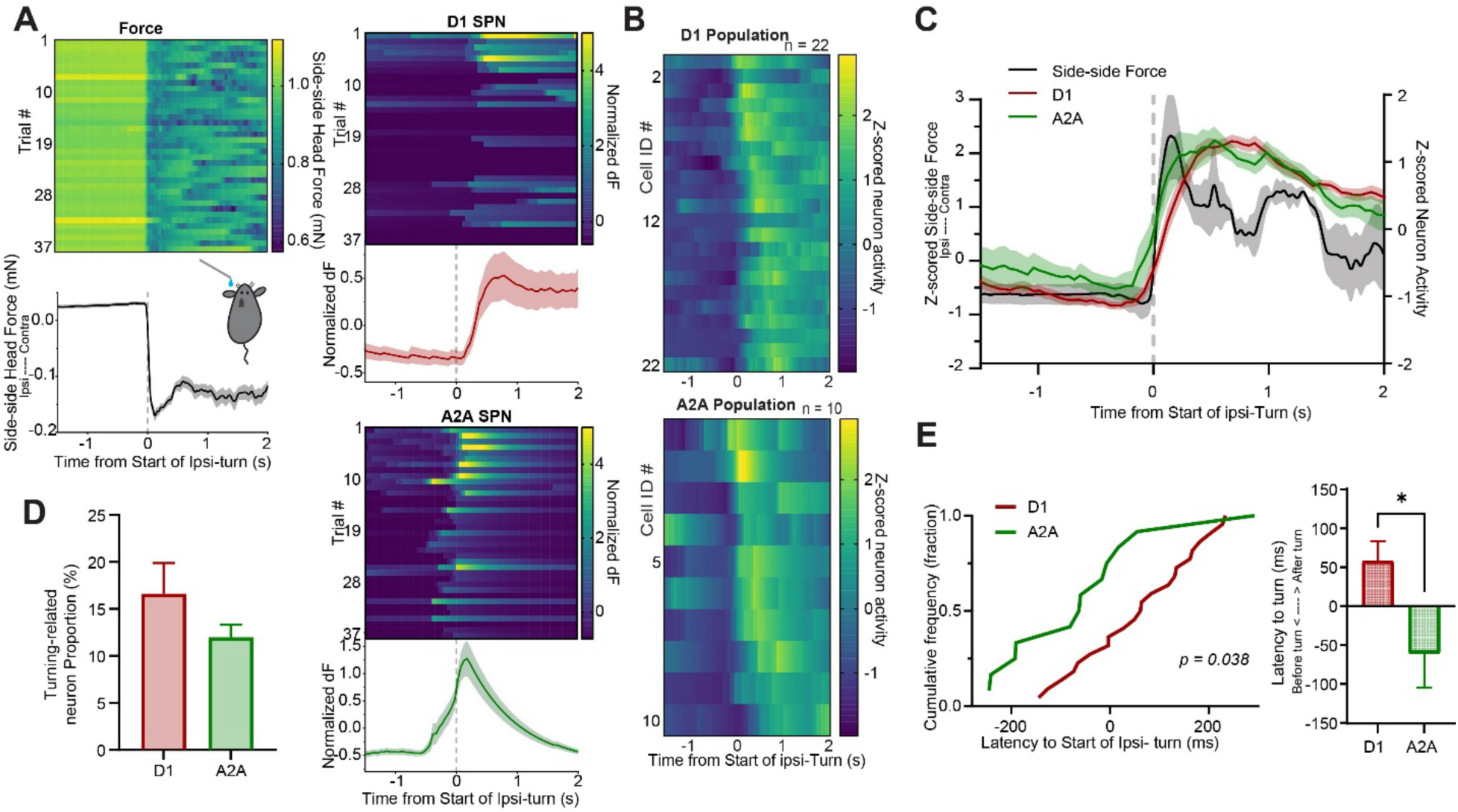
Coordination between striatal dSPNs and iSPNs during ipsiversive force exertion in head-fixed mice. A) Representative session showing ipsiversive force exertion (Left), with corresponding activity of example D1 SPNs (Top right) and A2A SPNs (Bottom right). B) Average calcium signals of all ipsiversive pushing-related dSPNs (Top, n=2, 22 cells) and iSPNs (Bottom, n=2, 10 cells), aligned to the start of force exertion and sorted by peak activity time (We recorded a total of 130 dSPNs and 94 iSPNs from 2 D1-Cre and 2 A2A-Cre mice. Specifically, A2A-Cre mice contributed 117 dSPNs and 32 iSPNs, whereas D1-Cre mice contributed 13 dSPNs and 62 iSPNs). C) Z-score of dSPNs activity (red, right y-axis),iSPNs activity (green, right y-axis) and lateral force (black, left y-axis) aligned to the start of force exertion. The SPNs are the same populations as shown in B). D) Proportion of ipsiversive pushing-related neurons. E) Latency (time lag between neural activity and behavioral measure) to the start of force exertion for dSPNs (red) and iSPNs (green). (Left) Cumulative distribution plots. A two-sample Kolmogorov–Smirnov test revealed a significant difference between the two distributions (p =0.038). Right, average latency to start of behavior. * p = 0.0147, two-tailed unpaired t-test.

## Discussion

In this study, we developed and validated a dual-color miniscope that enables simultaneous calcium imaging from two genetically defined neuronal populations in freely moving mice. By separating excitation paths, emission filters, and CMOS sensors, this system allows concurrent acquisition from two dynamic fluorescent indicators while minimizing optical crosstalk. We tested our system in the striatum, where dSPNs and iSPNs are spatially intermingled and hard to distinguish using single-channel imaging^29^. Dual-color calcium imaging therefore provides a uniquely powerful tool for studying neural activity in these two pathways under identical physiological and behavioral conditions. We used Cre-on and Cre-off AAV vectors to express two distinct calcium indicators in dSPNs and iSPNs ^30^, and two different LEDs in the excitation light path to excite GCaMP6s and jRCaMP1b.

Using this system, we observed consistent differences in the temporal relationship between dSPN and iSPN activity during goal-directed movements. Across multiple behavioral paradigms, dSPN activity usually preceded iSPN activity during contraversive movements, whereas this temporal relationship was reversed during ipsiversive movements. These timing differences were reproducible across different behavioral tasks and measurements. Our observations suggest that the relative timing of dSPN and iSPN recruitment is systematically related to movement direction and behavioral demands.

An influential finding is that, during contraversive turning, dSPNs and iSPNs are concurrently active^21^. However, we found that dSPN activity leads iSPN activity. Although dSPN activity preceded iSPN activity by 100-300 ms during contraversive turning, this difference could be very small when the velocity is low, which may explain why this difference in timing was not clearly observed in previous studies. Another possible reason previous work did not detect this timing difference is the limitation in the spatial resolution of the techniques used. For example, using dual-color photometry, Meng et al showed that dSPNs and iSPNs were co-activated during contraversive turning^11^. However, activity recorded using fiber photometry reflects bulk calcium signals in the FOV. These measurements may reflect the population average of the activity of stratonigral and striatopallidal neurons, or even the activity in non-neuronal cells. They lack sufficient temporal and spatial resolution to determine whether individual dSPNs and iSPNs are concurrently active ^31^.

Recent work showed that sensorimotor SPN activity is often highly correlated with movement velocity^32,33^. Studies have shown that activation of dSPNs alters movement velocity^34^, yet it remains unclear how the activity of dSPNS and iSPNs could be related to kinematics, as the timing and magnitude of activity in these two pathways have not been directly compared in the same animals. Using the switching task in freely moving mice and force measurements in head-fixed mice, we found that many dSPNs (∼40%) were activated during the acceleration phase, while many iSPNs (∼30%) were associated with deceleration. The distinct temporal dynamics of integration and decay can account for the direction-dependent timing of dSPN and iSPN activity, consistent with a leaky integrator model in which the direct pathway in a given hemisphere implements the inflow whereas the indirect pathway implements the leak^14^.

### Comparisons with previous systems

One approach to two-color calcium imaging is to use a single CMOS sensor with alternating excitation of two indicators ^38,39^. For example, the Inscopix nVue system collects all emitted fluorescence through a single emission path onto a single CMOS sensor. When the acquisition frame rate is set to 60 Hz, the effective sampling rate per channel is halved to 30 Hz. For dual-color imaging using GCaMP and RCaMP, this temporal splitting imposes a significant bottleneck: the RCaMP1b signal is typically too weak to be reliably captured at such high speeds. Consequently, the reduced exposure time per channel fails to accumulate sufficient photons, resulting in a signal-to-noise ratio (SNR) inadequate for tracking fast calcium dynamics. In contrast, our dual-CMOS design facilitates truly independent, simultaneous two-channel imaging. By eliminating the need for interleaved LED switching, both channels can be recorded at their full temporal resolution, ensuring stable and high-fidelity acquisition of dynamic signals from two distinct neuronal populations.

Our system uses separate CMOS sensors and emission filters for each channel (Figure 2). Each CMOS sensor is independently adjustable along the optical axis, allowing precise focal control for each channel. This compensates for chromatic aberrations introduced by the GRIN lens, addressing focal mismatches between fluorophores. In addition, the modular optical architecture supports interchangeable emission filters and dichroic mirrors, making the system compatible with a wide range of calcium indicators. Together, these features provide a flexible, scalable, and cost-effective platform for dual-color imaging in applications where signal independence and temporal fidelity are critical.

Recently, Dong et al. developed a dual-channel miniscope using a single LED excitation source to simultaneously capture dynamic GCaMP signals alongside a static dTomato reference^40^. Our design incorporates two independent LED paths with distinct wavelengths, enabling concurrent imaging of two dynamic signals from two distinct neuronal populations, rather than one dynamic and one static signal. With our dual-color miniscope, we achieved stable RCaMP signal acquisition even under GCaMP’s blue-light excitation, which typically accelerates RCaMP photobleaching^41^. This capability facilitates stable imaging of two dynamic channels, making it possible for future applications such as combining calcium indicators with neurotransmitter sensors or studying neuron–glia interactions in vivo.

## Conclusions

Simultaneous recording from multiple genetically defined populations within the same animal offers clear advantages for controlling behavioral variability and comparing relative activity patterns. At the same time, careful validation of optical crosstalk, sensor kinetics, and temporal alignment is essential when interpreting fine timing differences. As indicator technologies and imaging hardware continue to improve, combining dual-color imaging with faster reporters or direct electrophysiological approaches will be crucial for refining circuit-level models of basal ganglia function.

In summary, we present a dual-color miniscope that enables robust, simultaneous imaging of two neuronal populations in freely moving mice. Applying this tool to the study of striatal circuit dynamics reveals consistent, behavior-dependent differences in the relative timing of dSPN and iSPN activity across behaviors. These results shed light on how opponent basal ganglia pathways coordinate movement dynamics in real time, and demonstrate the utility of the dual-color miniscope in investigating temporal dynamics of different neuronal populations.

## Limitations of study

Several caveats should be noted, however, as they constrain the strength of conclusions that can be drawn about timing differences between dSPNs and iSPNs. First, the two calcium indicators used here (GCaMP6s and jRCaMP1b) exhibit distinct kinetics, including differences in rise time and decay. Such differences can influence apparent onset latencies, particularly when comparing signals across channels. Second, although our dual-CMOS architecture eliminates temporal interleaving of excitation, calcium imaging inherently provides indirect and temporally filtered measurements of neural activity. Accordingly, the observed difference between dSPN and iSPN activity can only be a rough estimate of the temporal lag in firing rates of these neuronal populations. Future studies employing faster indicators, voltage sensors, or electrophysiological recordings will be essential for resolving pathway dynamics at higher temporal resolution.

The direction-dependent ordering of dSPN and iSPN activity aligns with prior work linking striatal output to movement velocity and force generation, and suggests that relative pathway timing may reflect dynamic engagement of drive- and braking-related processes during action execution^13,14,35–37^. However, our data do not imply that dSPNs and iSPNs always operate in strict temporal segregation. For example, concurrent activity may be observed during behavioral conflict.

## Author Contributions

H. H. Y. conceived the concept of dual-color calcium recording system. J.Z., N.K. and H.H.Y. designed the experiments. J.Z. designed, assembled, and calibrated the circuit board and dual-color Miniscope. J.Z and J.K. performed testing in mice. J.Z, K.B. and F.H. analyzed data. J.K., K.B. and J.K. performed surgeries. J.K., J.Z. performed histology and confocal imaging. J.Z., F.H. & H.H.Y. wrote the manuscript.

## STAR Methods

### Experimental model and study participant details

All experimental procedures were approved by the Animal Care and Use Committee at Duke University. Male *D1-*Cre mice (Jackson Labs: Drd^1tm2.1Stl^) and *A2A-Cre* (Adora2A^tm1Dyj/J^) mice were used. Mice were between 3–8 months of age, group-housed under a 12:12 h light/dark cycle and tested during the light phase. All testing was performed during the light phase. Mice had ad libitum access to food and water, except during water-restriction training sessions. Only males were used; therefore, sex-based analyses were not performed.

### Key resources table

| REAGENT or RESOURCE | SOURCE | IDENTIFIER |
| --- | --- | --- |
| Bacterial and virus strains |  |  |
| AAV1-CAG-Flex-jRCaMP1b | Addgene | #100849-AAV1 |
| AAV9-CBA-DO(FAS)-GCaMP6s | Duke viral core | N/A |
| Chemicals, peptides, and recombinant proteins |  |  |
| DAPI | Sigma-Aldrich | D9542 |
| Isoflurane | Henry Schein | 1182097 |
| Paraformaldehyde | Sigma-Aldrich | P6148-1KG |
| Experimental models: Organisms/strains |  |  |
| Mouse: *Drd1<sup>tm2.1Stl</sup>/J* | The Jackson Laboratory | # 025700 |
| Mouse: B6;129-*Adora2a<sup>tm1Dyj</sup>/J* | The Jackson Laboratory | # 010687 |
| Software and algorithms |  |  |
| Prism 10 | GraphPad | https://www.graphpad.com |
| NeuroExplorer 5.436 | Nex Technologies | http://www.neuroexplorer.com/downloads/page/ |
| Matlab R2022a | MathWorks | https://www.mathworks.com/products/matlab.html |
| Cerebus Central Suite | Blackrock Neurotech | https://blackrockneurotech.com/research/support/software/ |
| Cortex | Motion Analysis, CA | https://www.motionanalysis.com/software/cortex-software/ |
| Zen | Zeiss | https://www.zeiss.com/microscopy/us/products/software/zeiss-zen.html |

### Method details

#### Viral vectors

AAV1-CAG-Flex-jRCaMP1b is from Douglas Kim & GENIE Project (Addgene viral prep # 100849-AAV1; http://n2t.net/addgene:100849; RRID:Addgene 100849). AAV9-CBA-DO(FAS)-GCaMP6s plasmid is from Bernardo Sabatini, and the vector is made by Duke Viral core.^42^

#### Stereotactic surgery

Animals were initially anesthetized with 3% isoflurane and maintained at 1-2% during surgical procedures. Meloxicam was administered for analgesia at a dose of 2 mg/kg body weight. After removing the scalp to expose the skull, a craniotomy was performed over the dorsal striatum (AP: 0.0∼1.0 mm from Bregma, ML: 2.0 ∼ 2.7 mm). Pulled pipettes were then used to inject the virus using a Nanoject III injector (Drummond Scientific, USA). The first virus was injected (250 nL of pAAV.CAG.Flex.NES-jRCaMP1b.WPRE.SV40) at two sites (AP: +0.25, +0.75 mm from Bregma, ML: 2.5 mm) each with 5 depths (DV: −2.8∼ −2.0 mm), at a rate of 1 nL/sec. The second virus (250 nL of AAV9-CBA-DO(FAS)-GCaMP6s) was injected at the same coordinates. The injection pipette was left in place for three minutes after injection before being slowly retracted.

Following viral delivery, tissue above the injection site was aspirated. A gradient-index (GRIN) lens (1.8 mm x 4.3 mm, Edmund Optics) was implanted in the dorsal striatum directly above the injection site (Supplementary Figure 6). Skull screws were placed in a square configuration around the surgical site, the lens was then secured to the skull using dental cement. Kwik-Sil was applied to protect the lens surface. Five to six weeks after GRIN lens implantation, base plating was performed under visual guidance of calcium signals to identify the optimal field of view (FOV).

#### Histology

After the experiments, mice were transcardially perfused with 0.1 M phosphate-buffered saline (PBS) followed by 4% paraformaldehyde (PFA) to confirm lens placement and viral expression. The mouse brains were then transferred to a 30 % sucrose solution and sliced coronally using a cryostat (Leica CM1850) and mounted with DAPI-containing mounting medium (Sigma-Aldrich) to visualize cell nuclei. Brain slices were imaged using either an inverted confocal microscope (Zeiss LSM 780) or an upright epifluorescence microscope (Axio Imager.M1, Zeiss).

#### Dual-color miniscope design: excitation path and fluorescence field alignment

To ensure effective and balanced excitation of both indicators, the blue (450–490 nm) and lime-green (547–572 nm) LED illumination fields were designed to be comparable in size and aligned along a common optical axis. A plano-convex lens in the excitation path was used to concentrate sufficient energy for reliable indicator excitation. To minimize optical aberrations while reducing miniscope weight, the excitation dichroic mirror (T510lpxr) was raised by 0.25 mm rather than incorporating an achromatic lens. Optical simulations performed in TracePro confirmed that the blue and lime-green illumination areas beneath the GRIN lens were well aligned. This alignment was further validated by imaging an optical resolution target with the assembled dual-color miniscope, which demonstrated precise overlap of the excitation fields (Supplementary Figure 2).

Due to its broader emission spectrum and greater distance from the excitation lens (2 mm), the lime-green LED required higher power than the blue LED to achieve equivalent excitation. Simulations using equal input power showed that the blue LED produced approximately 10-fold higher irradiance than the lime-green LED. To ensure equal excitation intensity (2 mW/mm²) beneath the GRIN lens for both LEDs, we used a higher-capacity power supply with parallel current sources to drive the lime-green LED.

#### Dual-color miniscope design: filter configuration and LED control

Each LED was paired with a dedicated excitation filter: a blue excitation filter (ET470/40x) for the blue LED and a lime excitation filter (ET560/25x) for the lime-green LED. The two excitation beams were combined using a dichroic mirror (T510lpxr) and directed into the GRIN lens via a main excitation dichroic mirror (69013bs).

In the emission path, fluorescence signals were filtered and directed to two separate CMOS sensors. The GCaMP6s channel used a green emission filter (ET525/36m), and the jRCaMP1b channel used a red emission filter (ET630/75m). A narrow-band emission filter was selected for the GCaMP6s channel to reduce background fluorescence. An emission dichroic mirror (T550lpxrxt) splits the fluorescence and directs it to the corresponding CMOS sensors.

#### Calcium imaging acquisition

All experimental devices were synchronized using a common trigger signal initiated by Computer 1. Computer 1 first generated a start pulse that was sent to Miniscope 1, which in turn produced a rising-edge electronic signal distributed to all other connected devices, including Miniscope 2 and the behavioral tracking system. This ensured precise temporal synchronization between the two imaging channels and behavioral recordings (Supplementary Figure 1).

Calcium imaging was captured by a CMOS imaging sensor connected to a data acquisition (DAQ) system and a USB host controller. Images were acquired at 20 frames per second and saved as uncompressed “.avi” files through the DAQ software.

Because jRCaMP1b fluorescence is weaker than GCaMP6s due to competition between the two viral vectors expressed in the same region, we enhanced the jRCaMP1b video data prior to signal extraction. The enhancement procedure consisted of two steps: contrast enhancement and image sharpening. The results of this processing are illustrated in Supplementary Video 2.

In the enhancement function, we first performed contrast stretching around a midpoint of 128, followed by multiplicative brightness scaling, and finally clipped each channel to the range [0, 255]. The corresponding equations are as follows:

1. Contrast factor conversion (standard contrast slider formula):

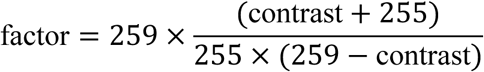 where *contrast* is a user-defined parameter.
2. Contrast adjustment:

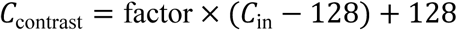 where *C*_in_represents the input image data.
3. **Brightness adjustment:**

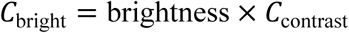 where *brightness* is another user-defined parameter.
4. **Output combination:**

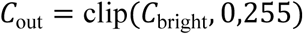

Finally, image sharpening equivalent to the MATLAB *imsharpen* function was applied to improve edge definition and enhance the visibility of neuronal structures.

To ensure the temporal fidelity of calcium imaging data, we quantified frame drops across all recording sessions using a dual-timestamp synchronization system. Hardware-generated frame flip signals, which reflect the actual timing of frame acquisition, were recorded independently of software timestamps that indicate when frames were written to disk. By comparing these two timestamps, we precisely identified the occurrence and timing of dropped frames. Across 52 calcium imaging recordings included in the biological analyses, only 8 exhibited any frame loss. Of these, 7 recordings dropped fewer than 15 frames over the entire session (typically ∼3,600 frames acquired at 20 Hz over 3 min). One recording initially showed 158 dropped frames; however, nearly all losses occurred within the first 5–10 s of acquisition. To maintain analytical rigor, this initial segment was excluded from subsequent analyses. Importantly, none of the recordings used for temporal analyses (including estimation of the ∼110 ms dSPN–iSPN lag) exhibited more than 15 dropped frames. Together, this dual-timestamp validation approach ensures that temporal measurements remain accurate and robust despite occasional frame loss.

#### The target switching task

The switching task was designed to elicit reliable turning behavior in mice (Figure 4). Mice moved between two water spouts positioned 50 mm apart. An LED positioned above each spout served as a visual cue. When it is turned on, it indicates that the spout next to the LED has water. Thus the mouse has to switch between different target spouts to obtain water, and in this process exhibit both ipsiversive and contraversive turning behavior. Mouse behavior was tracked using 3D motion capture^34,36^. Two infrared reflective markers (6.35 mm diameter), spaced 2 cm apart, were mounted to the dual-color miniscope to track mouse head movements. Seven Raptor-H Digital Cameras (Motion Analysis, CA; 100 Hz sampling rate) captured the positions of the markers, which were converted to Cartesian coordinates (x, y, z) using Cortex software (Motion Analysis, CA). MATLAB interfaced with Cortex in real time to control reward delivery and LED switching during the task. A reward (∼12 µL of 20% sucrose) was delivered every 600 ms. Prior to imaging, mice were trained for 2–4 hours over the course of one week to perform immediate turning responses following LED cue switching. An immediate turn was defined as a rapid head movement from one target location to the other within 2 seconds of the LED onset. Trials in which mice failed to turn immediately included behaviors such as grooming, rearing, orienting, or exploratory scanning. In some cases, animals made repeated left–right head movements before completing the turn, which was also classified as a non-immediate response. Training continued until each mouse achieved at least 80% immediate turns during a 10-minute session.

#### Head-fixation system with continuous load cell force monitoring

The head-fixation system uses three load cells, mounted to a plastic frame, to measure forces generated by mouse head movements along three orthogonal directions. A steel head bar, implanted during surgery with dental cement, was attached to the frame using small pinch clamps during experiment (**Figure 5A**). The analog voltage signals from the load cells were sampled at 10 kHz using a data acquisition system (Blackrock Neurotech) and calibrated to force values for subsequent analysis^25,26,43^.

At the beginning of training, the spout was positioned close to the mouse within easy reach. Once the mouse adapted to the current distance, the spout was gradually moved farther away until the mouse had to exert substantial force (load cell voltage exceeding 500 mV) toward the spout to get the reward. Mice were trained until they get the reward consistently (2-4h of training over 1 week).

### Quantification and statistical analyses

#### Behavioral analysis

For the target switching task, two infrared reflective head markers were used for tracking mouse and target movements. These markers were captured by eight Raptor-H Digital Cameras (Motion analysis, CA, 100 Hz sampling rate). The data was transformed into Cartesian coordinates (*x*, *y*, and *z*) by the Cortex program (Motion Analysis, CA). MATLAB communicated with the Cortex program (Motion Analysis) online to control reward delivery based on mouse behavior. The positions of the two spouts were also tracked and used to define the left–right axis of movement. The known distance between the spouts served as a reference to convert motion capture coordinates into real-world distances. Head movements were quantified as changes in position along this axis, and head velocity was calculated as the temporal derivative of head position.

In the switching task, the “Head Velocity Peak” event was defined as the local maximum of the head position signal along the left–right axis. In the load cell experiments, the “Start of Turn” event was defined as the threshold-based onset of increases in the continuous side-to-side force signal. Head turning was captured by load-cell voltage, and calibration force-voltage was used to transfer the voltage to force. For each event type, peri-event behavioral data from −1.5 s to +2 s were extracted, averaged, and z-scored across animals.

#### Neural signal extraction

Calcium imaging videos were imported into MATLAB for non-rigid motion correction analysis^44^, followed by signal deconvolution with the constrained non-negative matrix factorization (CNMF-E) algorithm^45^. Seed pixels were initialized with a minimum local correlation value of 0.8 and a minimum peak-to-noise ratio of 10. To ensure spatial specificity, the minimum number of non-zero pixels per neuron was set to 10, based on the resolution of the CMOS sensor (1 µm/pixel).

Raw calcium traces were saved and, after alignment with behavioral timestamps, segmented into peri-event epochs based on the same behavioral events within the same recording. Turning-related SPNs were defined as those exhibiting activation during more than half of the turning events within a one-second window centered on the head velocity peak. For each turning-related cell, responses were averaged across trials and z-scored across neurons.

The SNR of each neuron is calculated from raw data after neuron activities extraction. To ensure the neuron activities are not from the noise background, we use the following equation to calculate the SNR:

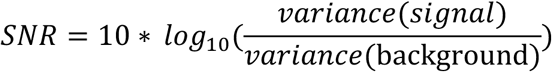

Normally, the selected neuron traces SNR is over 20, which illustrate the clearer identification of spike events and better separation from noise.

#### Statistical analysis

dSPNs and iSPNs from both D1-Cre and A2A-Cre mice were grouped together during the statistical analysis. All statistical analyses were performed in MATLAB and GraphPad Prism 7.0. Sample size was not determined *a priori*. All error bars represent SEM. Significance levels were set to P<0.05.

## Data and code availability

All data and Matlab codes used in the present study are available upon request.

## Ethical compliance

All animal experiments were conducted in accordance with the ARRIVE 2.0 guidelines to ensure transparent and reproducible reporting of in vivo research. Experimental design, sample size, randomization, and data analysis procedures were planned to minimize bias and ensure rigor. Details regarding animal housing, husbandry, anesthesia, analgesia, surgical procedures, and humane endpoints are described in the Methods section. All procedures were approved by the Duke University Institutional Animal Care and Use Committee (protocol A263-16-12) and were carried out in accordance with institutional and national ethical standards for the care and use of laboratory animals. All research reported in this study was approved by Duke University Institutional Animal Care and Use Committee (protocol A263-16-12).

## Acknowledgements

This research was supported by NS094754 to HHY.

