## Supplementary data for "A dual-color miniature endoscope for calcium imaging in behaving mice"

**Supplementary Information**

**Video S1:** Simultaneously record direct and indirect pathway neurons with open field behavior

**Video S2:** RCaMP channel enhancement

### Supplementary Figures

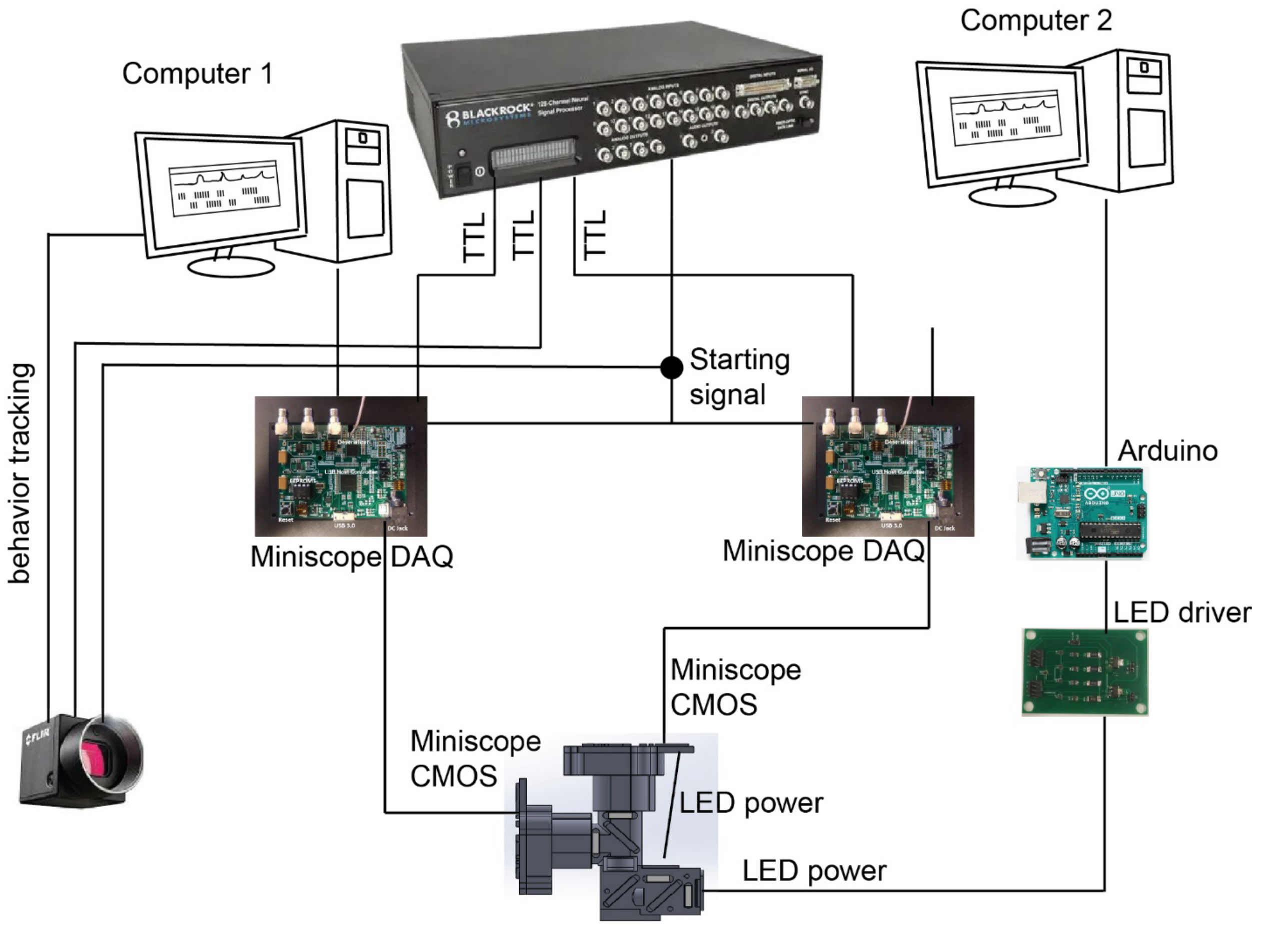


**Supplementary Figure 1. Dual-color miniscope system**

Computer 1 controlled the GCaMP6s channel DAQ and the behavioral tracking system. Computer 2 controlled the lime LED control and jRCaMP1b channel data collection. All signals were synchronized and recorded using a Blackrock data acquisition system.

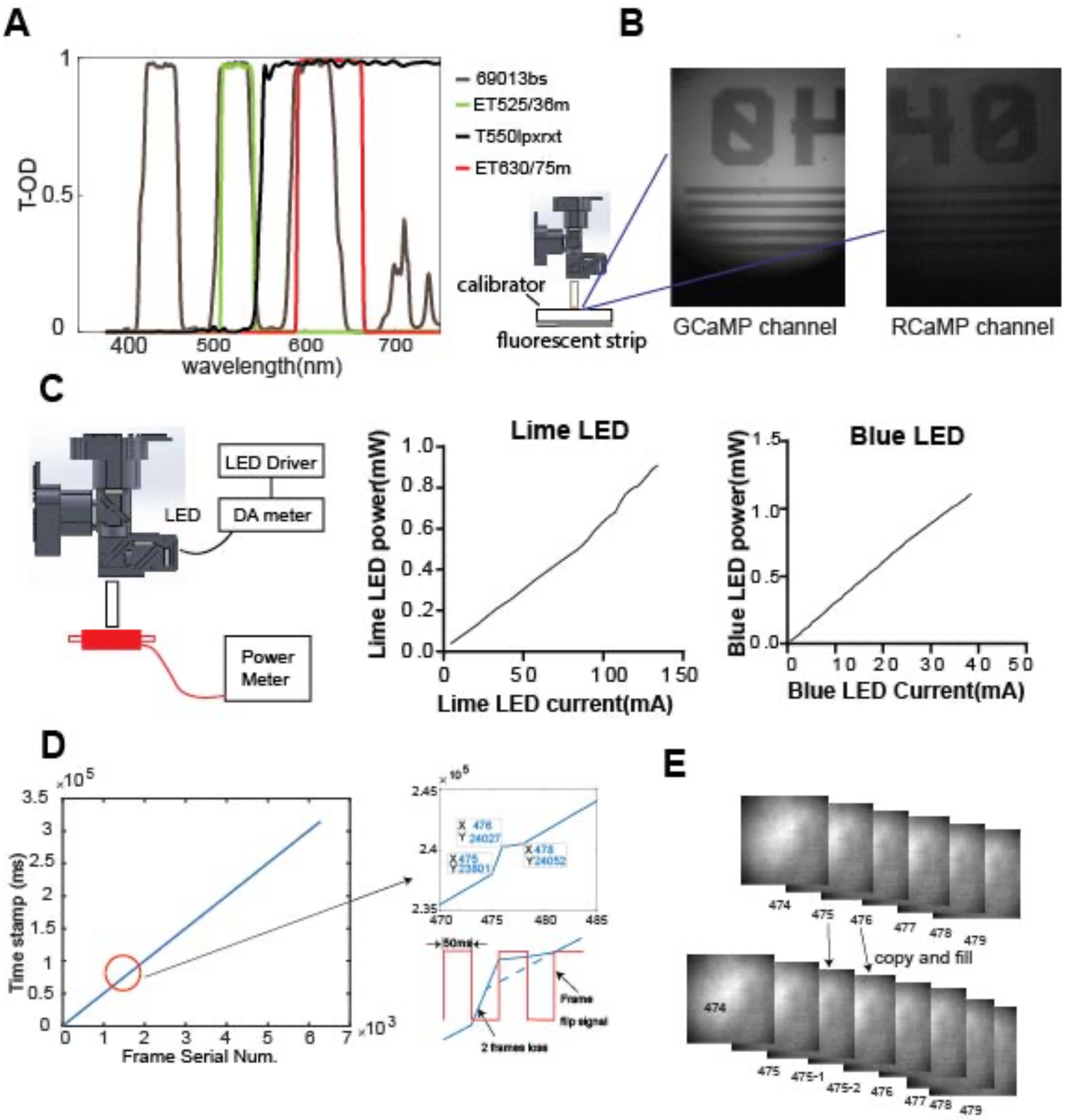


**Supplementary Figure 2. Performance of dual-color miniscope**

**A)** Primary filters and dichroic mirror wavelengths shows minimal spectral overlap between components.

**B)** Images captured by two different CMOS sensors using a fluorescence calibrator show good focus for each sensor.

**C)** LED output power was measured using a Thorlabs PM100D power meter positioned beneath the GRIN lens. The blue LED current was varied in 5 mA steps, while the lime LED current was adjusted in 10 mA steps.

**D)** Timestamp (ms) plotted against miniscope frame index. Deviations from linearity reveal dropped frames during acquisition.

**E)** Corrected frame sequence with missing frames replaced by duplication of preceding frames (Extremely rare scenario).

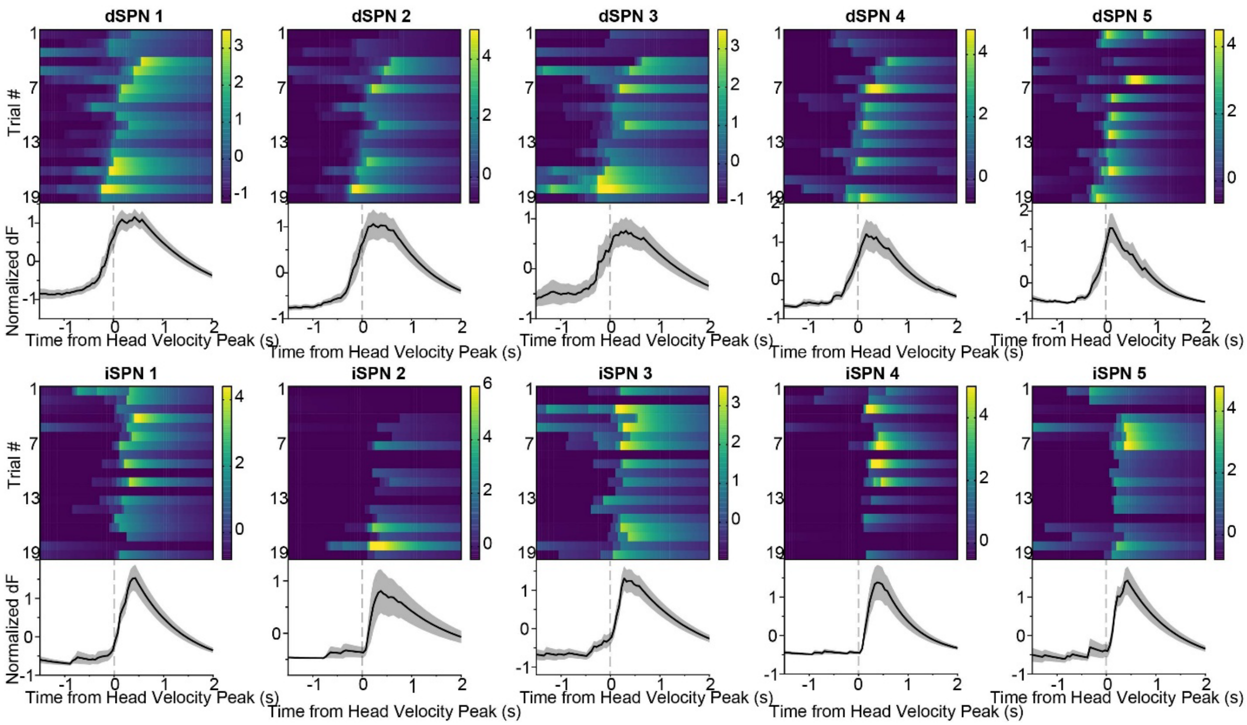


**Supplementary Figure 3. Representative neural activity during contraversive turning in the switching task.**

Representative dSPNs and iSPNs showing activity aligned to contraversive turning (velocity peak).

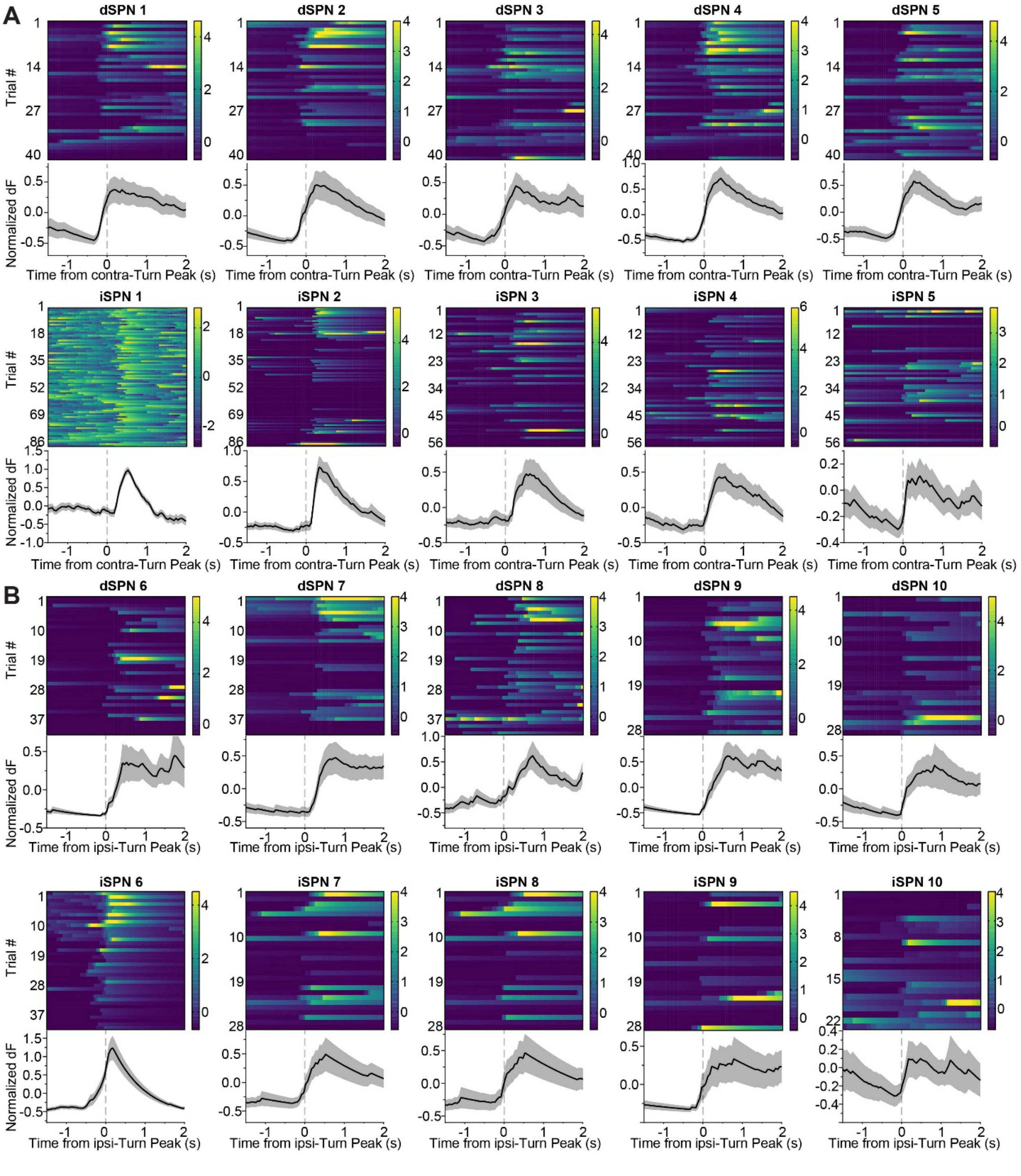


**Supplementary Figure 4. Representative neural activity during contraversive and ipsiversive force exertion in head-fixed mice**

A) Representative dSPNs and iSPNs aligned to contraversive force exertion.

B) Representative dSPNs and iSPNs aligned to ipsiversive force exertion.

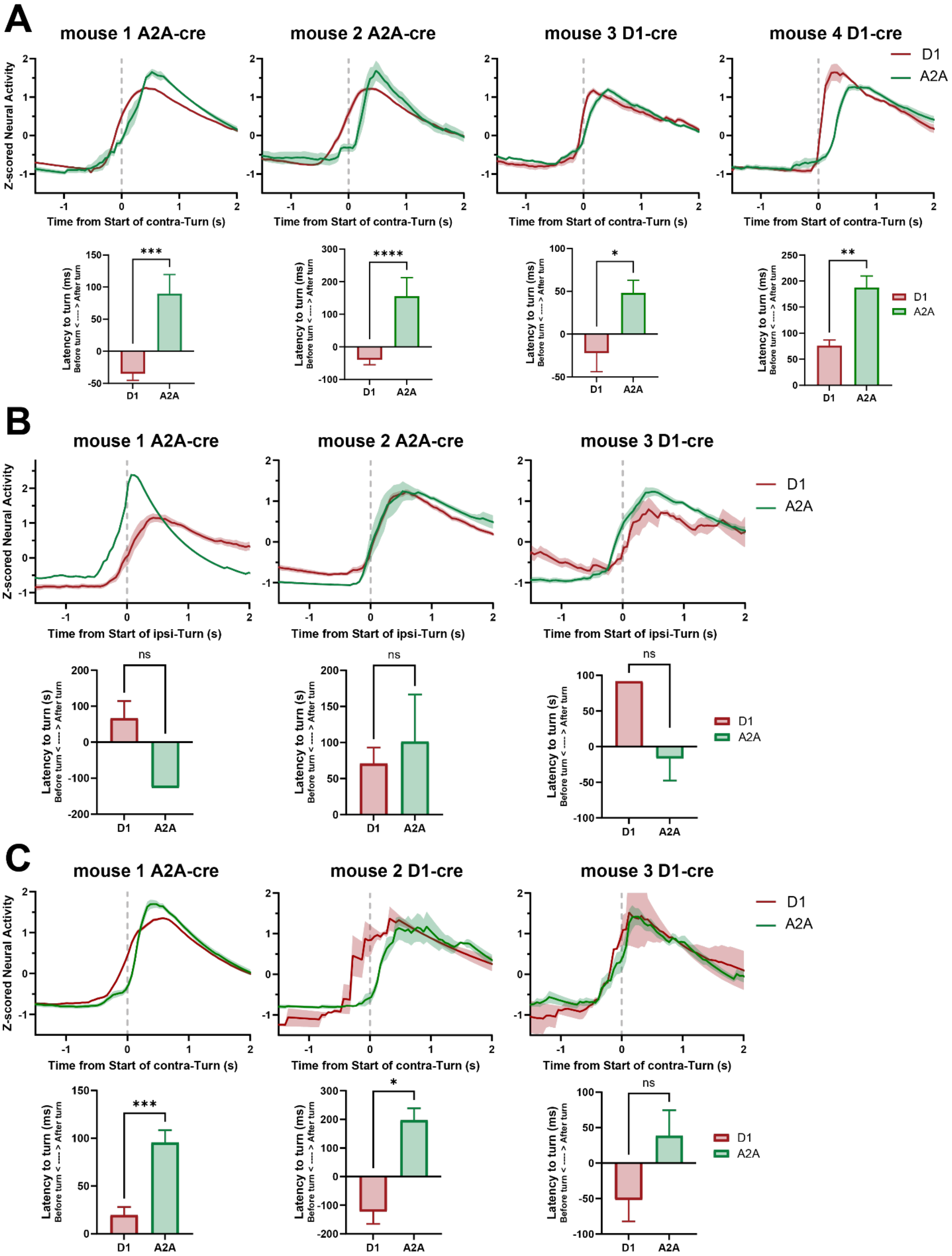
A
mouse 1 A2A-cre
mouse 2 A2A-cre
mouse 3 D1-cre
mouse 4 D1-cre
D1
A2A
Z-scored Neural Activity
Time from Start of contra-Turn (s)
Latency to turn (ms)
Before turn < ---- > After turn
***
****
*
**
B
mouse 1 A2A-cre
mouse 2 A2A-cre
mouse 3 D1-cre
Time from Start of ipsi-Turn (s)
Latency to turn (s)
ns
C
mouse 1 A2A-cre
mouse 2 D1-cre
mouse 3 D1-cre
Time from Start of contra-Turn (s)
Latency to turn (ms)
***
*
ns

**Supplementary Figure 5. dSPN and iSPN activity in individual mice.**

A) Activity of contraversive turn-related dSPNs (red) and iSPNs (green) in head-fixed mice. Bottom: Latency to head velocity peak for dSPNs and iSPNs during contraversive turns.

B) Same as (A), for ipsiversive turn-related neurons.

C) Same as (A), for contraversive turn-related neurons during the switching task. (**** $p < 0.0001$, *** $p < 0.001$, ** $p < 0.01$, * $p < 0.05$, ns, $p > 0.05$, two-tailed unpaired t-test).

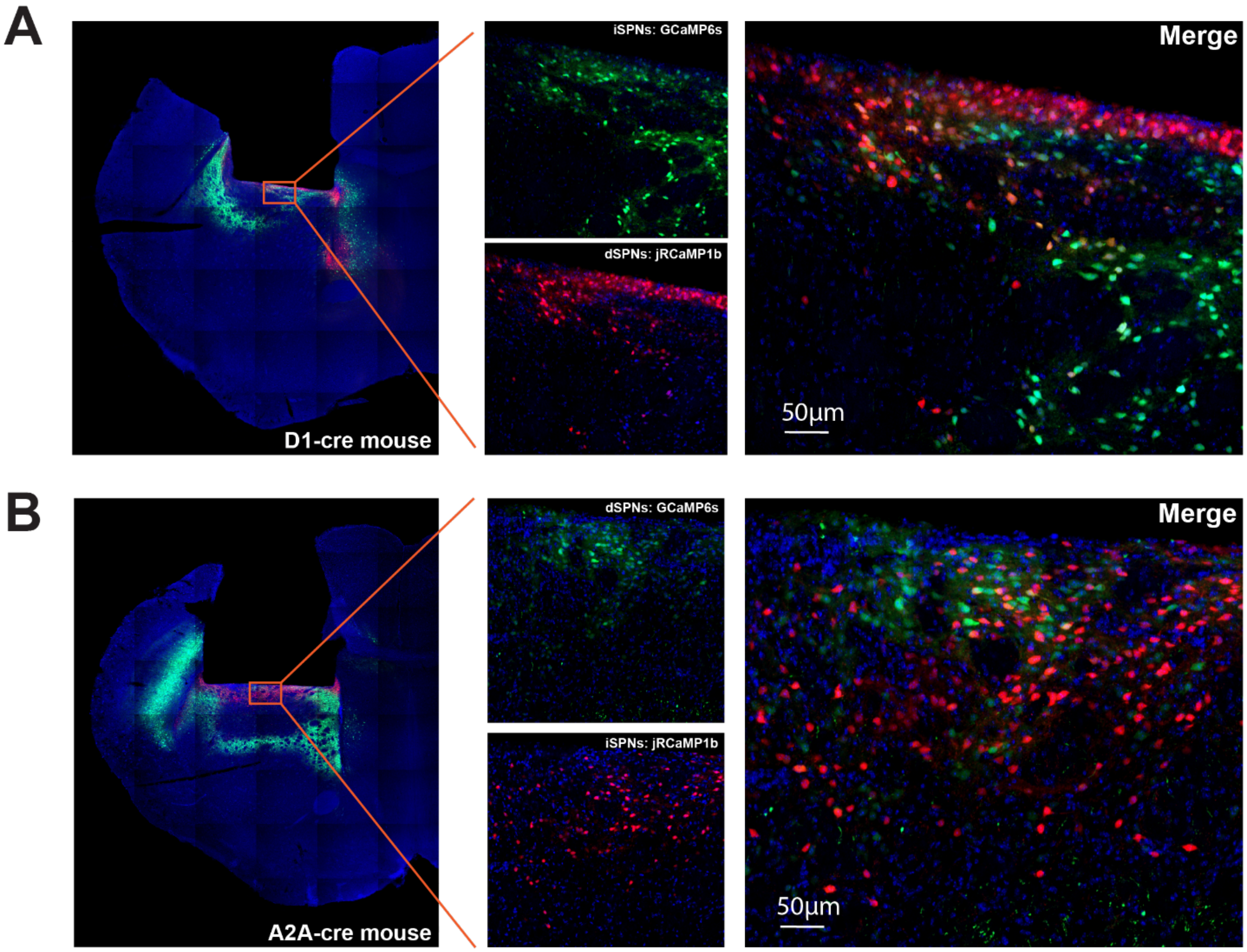


**Supplementary Figure 6. Histological verification of GCaMP6s and jRCaMP1b expression.**

A) Left, virus injection to the dorsal striatum of a D1-Cre mouse. Middle, dorsal striatum with GCaMP6s expression in iSPNs (Top) and jRCaMP1b expression in dSPNs (Bottom) Right, merged image showing non-overlapping expression of the two indicators. B) Histology from an A2A-Cre mouse, with dSPNs expressing GCaMP6s and iSPNs expressing jRCaMP1b in the dorsal striatum.
